# The Association between Tuberculosis and Diphtheria

**DOI:** 10.1101/219584

**Authors:** S. Coleman

## Abstract

This research investigates the now-forgotten relationship between diphtheria and tuberculosis. Historical medical reports from the 19^th^ century are reviewed followed by a statistical regression analysis of the relationship between the two diseases in the early 20^th^ century. Historical medical records show a consistent association between diphtheria and tuberculosis that can increase the likelihood and severity of either disease in a co-infection. The statistical analysis uses historical weekly public health data on reported cases in five American cities over a period of several years, finding a modest but statistically significant relationship between the two diseases. No current medical theory explains the association between diphtheria and tuberculosis. Alternative explanations are explored with a focus on how the diseases assimilate iron. In a co-infection, the effectiveness of tuberculosis at assimilating extracellular iron can lead to increased production of diphtheria toxin, worsening that disease, which may in turn exacerbate tuberculosis. Iron-dependent repressor genes connect both diseases.

## Introduction

The aims of this research are to resurrect long-forgotten knowledge about the relationship between tuberculosis and diphtheria and to support it with statistical analysis of historical data. This research began with an exploration of co-infections and associations between common infectious diseases in the United States using public health records. Among other findings, the analysis turned up a possible but unexpected correlation between tuberculosis and diphtheria. This was investigated subsequently in historical medical reports, as described below, revealing that, indeed, such a relationship was known to medicine in the late 19^th^ century. But knowledge of this seems to have waned as vaccination reduced the incidence of diphtheria. The historical relationship between diphtheria and tuberculosis is tested with modern statistical methods that were unavailable in that era. The statistical analysis focuses on the relationship between the two diseases in five American cities in the early 20^th^ century using weekly public health reports over a period of several years.

Medical authorities of the past did not have an explanation for the connection between tuberculosis and diphtheria, and this holds true today. Alternative explanations for the association are discussed here with a focus on how the two diseases assimilate iron in the body. Tuberculosis and diphtheria have a close phylogenetic relationship in the actinobacteria phylum, order corynebacteriales. As such, they share important commonalities in their cell biology and, specifically, in their iron-dependent repressor genes that uniquely control the assimilation of iron in both diseases.

## Method of Analysis

### Historical Review

The historical relationship between tuberculosis and diphtheria was investigated by an online search of historical medical texts, primarily from the late 19^th^ century. At that time both diseases were still very common and deadly, and the diphtheria bacterium had been identified in the 1880s by Klebs and Loeffler. The search was done in English, German and French through Google, Google Scholar and Google Book Search. Search terms were used to find medical accounts that referred to close or timely associations between the two diseases, such as, “with”, “in connection with”, “after”, “secondary infection’, and “mixed infection.”

A search of contemporary medical reports did not find any references to a causal or correlational association between the two diseases, but records from the late 19^th^ century tell a different story. Among German reports, an 1885 publication describes co-infections of diphtheria and tuberculosis among children with tuberculosis of bones and joints.[1] Diphtheria affected about 10 percent of such cases. A report from 1899 relates how tuberculosis can follow diphtheria and that diphtheria can infect persons with long-term tuberculosis infections.[2] Of 459 persons who died of diphtheria over an eight year period 95 (21%) had tuberculosis. In 37% of long-term tuberculosis cases there was a new eruption of tuberculosis with diphtheria, and almost one-third of children who had previously had tuberculosis got a new outbreak when infected with diphtheria. Additional research shows tuberculosis often appearing after diphtheria, or diphtheria occurring when there was primary tuberculosis of the intestines.[3] Of 714 sections taken from diphtheria patients who died between 1873 and 1894, tuberculosis was found 140 times (20%) in various organs.

A French study on the relationship between tuberculosis and diphtheria discusses these as family diseases, as they often occurred together in the same family.[4] The author asserts that neither directly causes the other, but each creates “favorable ground” for the other. A French analysis of children dying of diphtheria finds 41% with latent tuberculosis; diphtheria is aggravated by tuberculosis, while diphtheria also wakes up latent tuberculosis.[5] Similarly, a study of 150 children with tuberculosis in St. Petersburg reports that 39 (26%) had tuberculosis, affecting various organs.[6] The author concludes that the presence of tuberculosis predisposes to diphtheria, diminishes resistance, and worsens prognosis.

### Statistical Analysis

As the historical analysis reveals, at least 20% of diphtheria cases and possibly more involved co-infections with tuberculosis. The task of statistical analysis is to try to detect and estimate this association. Both tuberculosis and diphtheria are highly infectious airborne diseases. After infection, however, their pathogenesis and progression are very different. Tuberculosis was an endemic disease historically, while diphtheria had annual cycles. Tuberculosis is slow to develop and a person can have either an active case or a latent case, which can become active after a long delay. It affects lungs but also body organs and bones. Diphtheria has a short incubation period before it becomes fully established, and it mainly attacks the throat and tonsils. The damage from diphtheria throughout the body is caused by a toxin produced by a virus that has infected the bacterial DNA. The diseases have two aspects in common than can contribute to co-infections: both often attack children at young ages, and people can be asymptomatic carriers of either disease. These factors also foster the spread of infection within families.

This analysis relies on cases of diphtheria and tuberculosis reported to public health authorities in five American cities in the early 20^th^ century. These include the four largest American cities of the time—New York City, Chicago, Philadelphia, and Detroit—and Boston. This data is available from Project Tycho at the University of Pittsburgh.[7] This database was used previously to study the historical associations between measles and pertussis [8], and between varicella and scarlet fever.[9] Cases were reported on a weekly basis. Data on diphtheria extends from about 1916 to 1947 for most cities (from 1915 in Detroit) and is fairly complete back to 1907 in Philadelphia. Tuberculosis data runs from about 1906 to 1923. So the analysis for each city is based on the years when data on both diseases was available. An unresolved question is how cases might have been reported when both diphtheria and tuberculosis might have been involved. There is no designation for co-infections in the data. As with aggregate data generally, it is not possible to determine how often an association between the two diseases might involve the same person, the same person at different times, or whether it may represent an association at the family or community level. Note also that there is an inherent delay between the time a disease is diagnosed and when it appears in a public health report.

The relationship between diphtheria and tuberculosis is estimated with an ordinary least squares (OLS) regression model of weekly data for each city over the entire span of years with data. Diphtheria cases are considered the dependent variable, as it is more likely that among individuals tuberculosis preceded a diphtheria infection, although there may be a reciprocal effect. The weekly diphtheria data have a strong autocorrelation across time, which can cause problems for a regression analysis. The estimated coefficient for tuberculosis in the regression model is not affected, but the estimate of the standard error in the coefficient is likely to be underestimated, making the results more certain then deserved. To avoid this problem, OLS analysis was done using Gretl, an econometric software package that estimates a robust standard error, correcting any heteroskedasticity and autocorrelation problems in the standard errors.[10]

Because the number of diphtheria cases varies during the year, usually being highest in the winter and lowest in late summer, the regression model includes polynomial terms to the fourth degree as a function of week to model the annual cycle. (Higher order polynomial terms were not statistically significant.) Sometimes a sinusoidal model is used to estimate diseases with annual cycles, but it is not the best approach here. The peaks in the diphtheria cycles are too steep for a sinusoidal model, and the number of diphtheria cases varies substantially from year to year. (There are no weeks with zero diphtheria cases, which might otherwise be a problem for the regression model.) Because of the strong yearly variation in diphtheria rates, dummy variables are included in the model to annual changes. The final independent variable is the number of weekly tuberculosis cases. Because of the large number of coefficients in the model, only the coefficient for tuberculosis is reported along with summary information about the model. The estimated model for weekly diphtheria and tuberculosis case data is

Diphtheria cases = constant + b_1_ Tuberculosis cases + b_2_ week + b_3_ week^2^ + b_4_ week^3^ +b_5_ week^4^ + dummy variables for each year (0 or 1)

## Results

The five cities vary from year to year in the number of tuberculosis and diphtheria cases, as well as in weekly totals. Data on total reported cases for each city in each year from 1917 to 1923 (from 1916 for Detroit) are in Tables 1-5. Chicago (Figure 1) shows a typical pattern across years, with tuberculosis cases exceeding diphtheria. But tuberculosis rates were in general decline in the last few years of data. This yearly pattern is similar to the other cities except Boston and Detroit where diphtheria cases exceeded tuberculosis in some years. The reasons for such variations over time and across cities are unknown. The trend in weekly cases within a year, averaging across all the years, is illustrated in Figure 2 for Chicago. Diphtheria has a strong annual cycle while tuberculosis cases are relatively steady from week to week.

**Figure 1.**
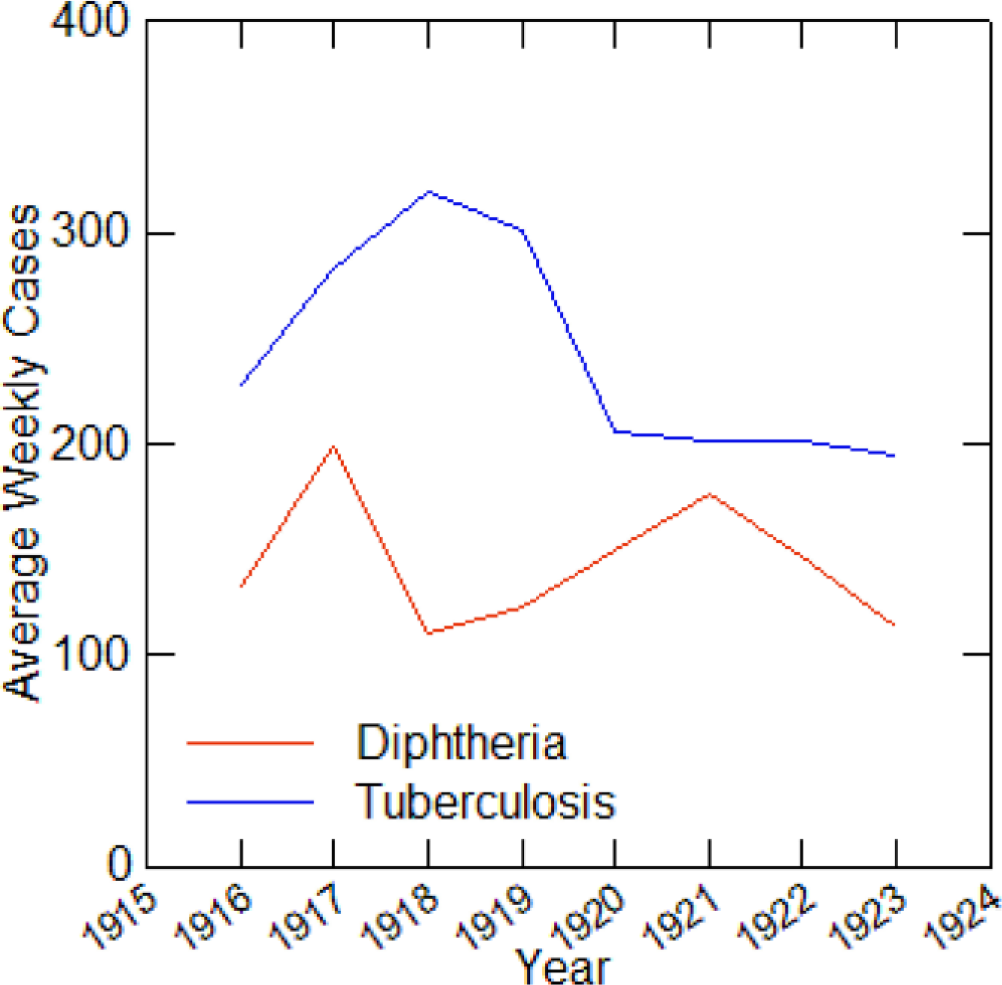
Average weekly diphtheria and tuberculosis cases for each year in Chicago, 1916-1923.

**Figure 2.**
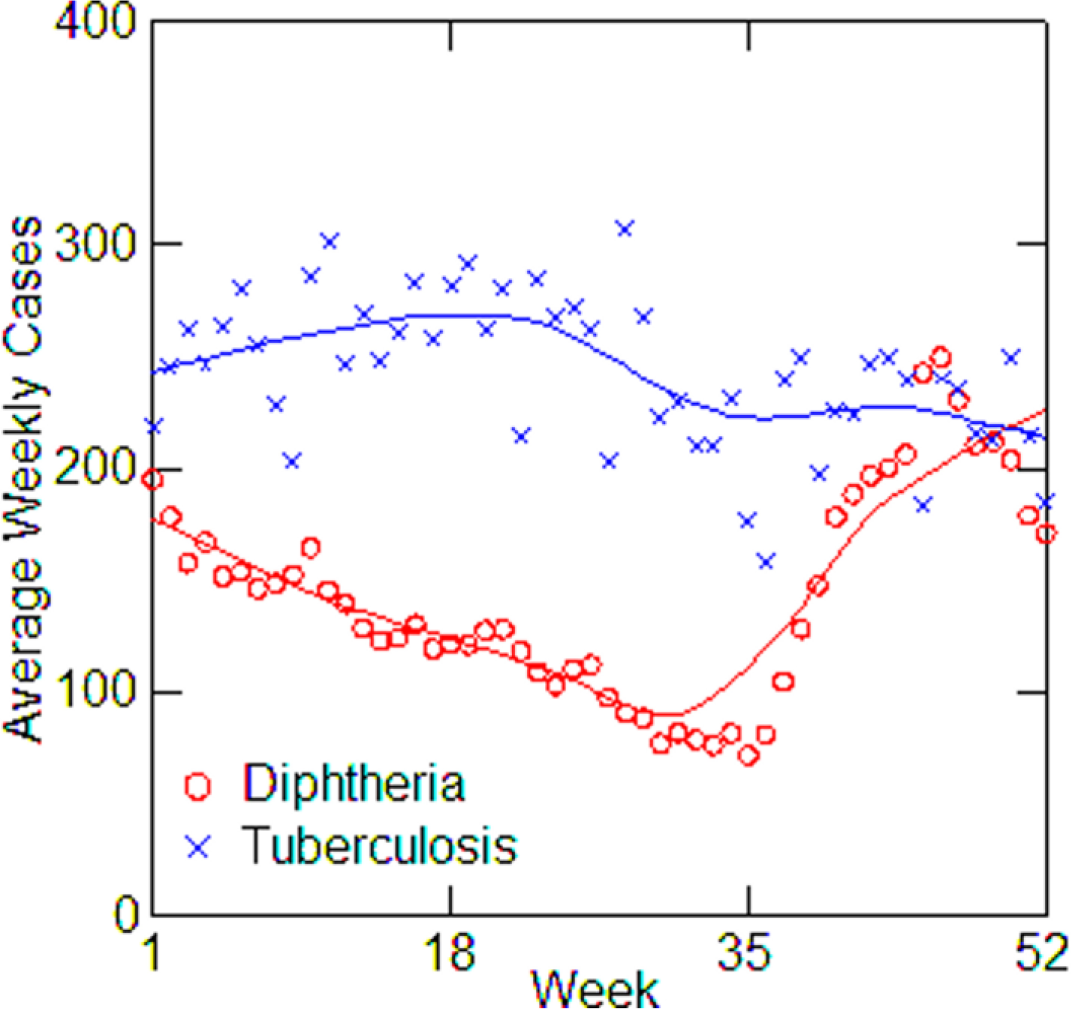
Average diphtheria and tuberculosis cases for each week of the year in Chicago, averaging over all years from 1916 to 1923, with LOWESS smoothing.

**Table 1.**
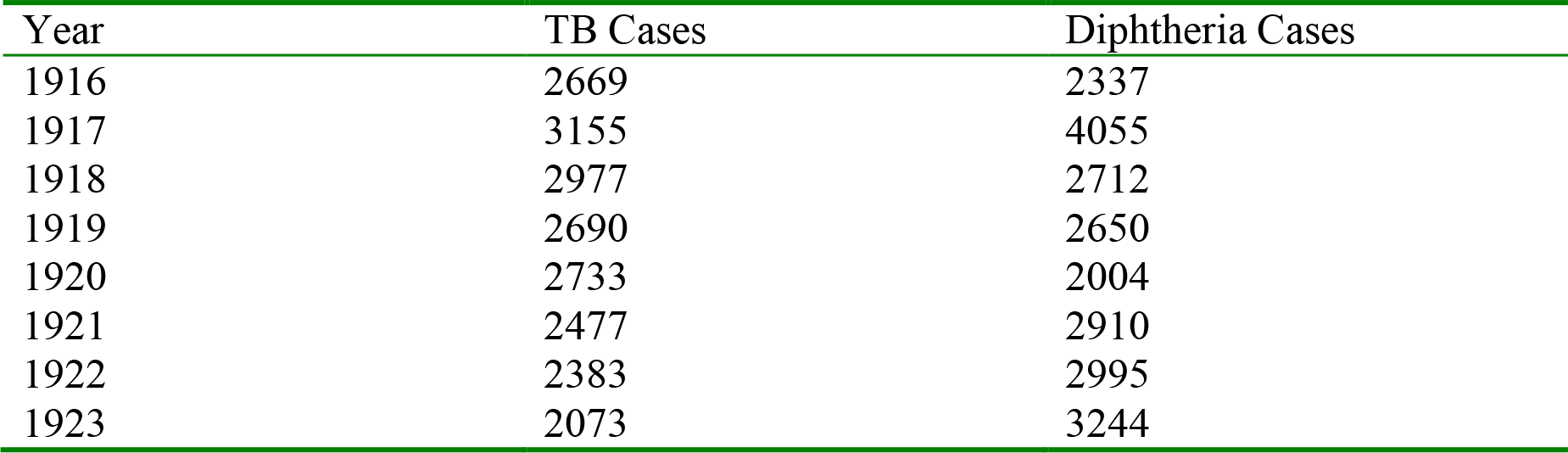
Boston, TB and diphtheria cases by year, 1916-1923.

**Table 2.**
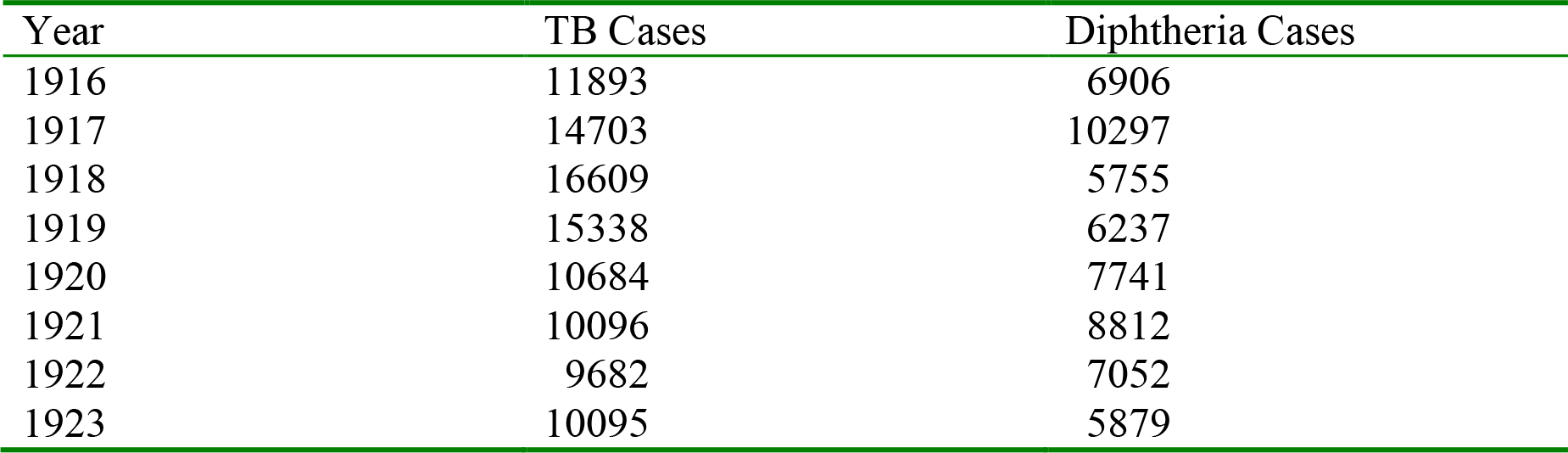
Chicago, TB and diphtheria cases by year, 1916-1923.

**Table 3.**
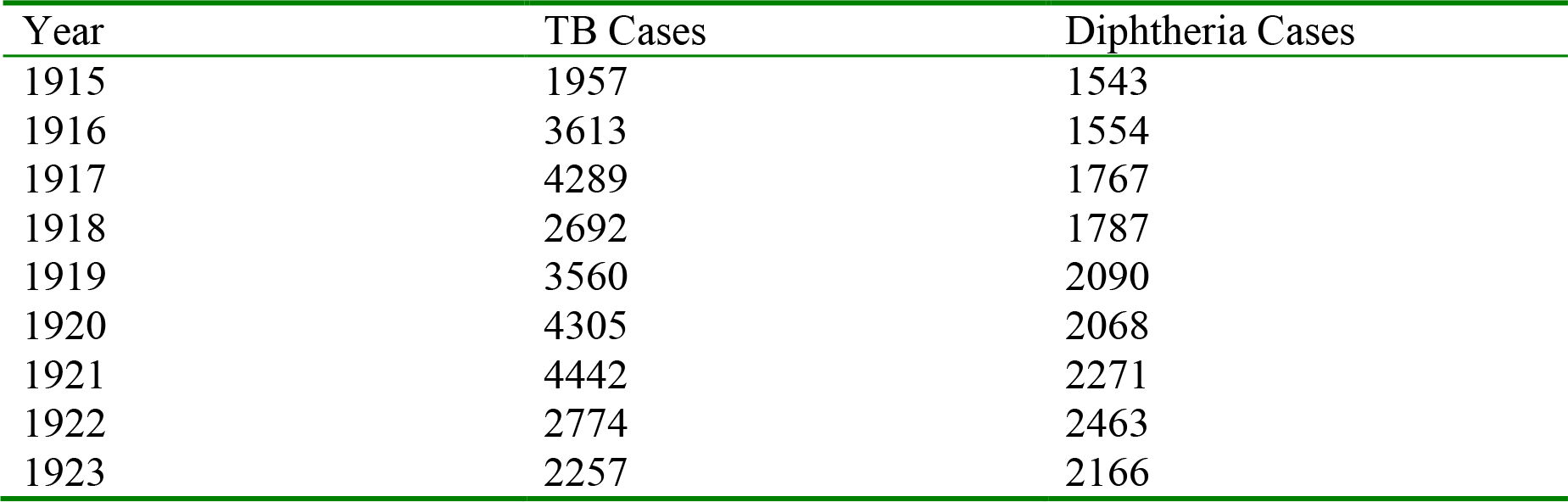
Detroit, TB and diphtheria cases by year, 1915-1923.

**Table 4.**
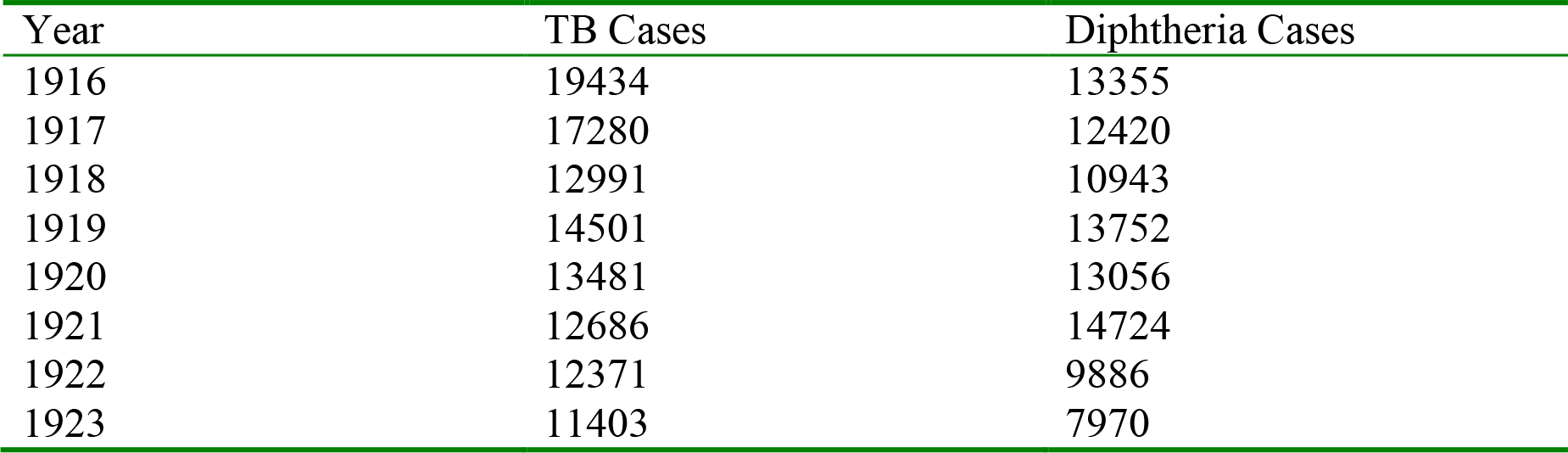
New York City, TB and diphtheria cases by year, 1916-1923.

**Table 5.**
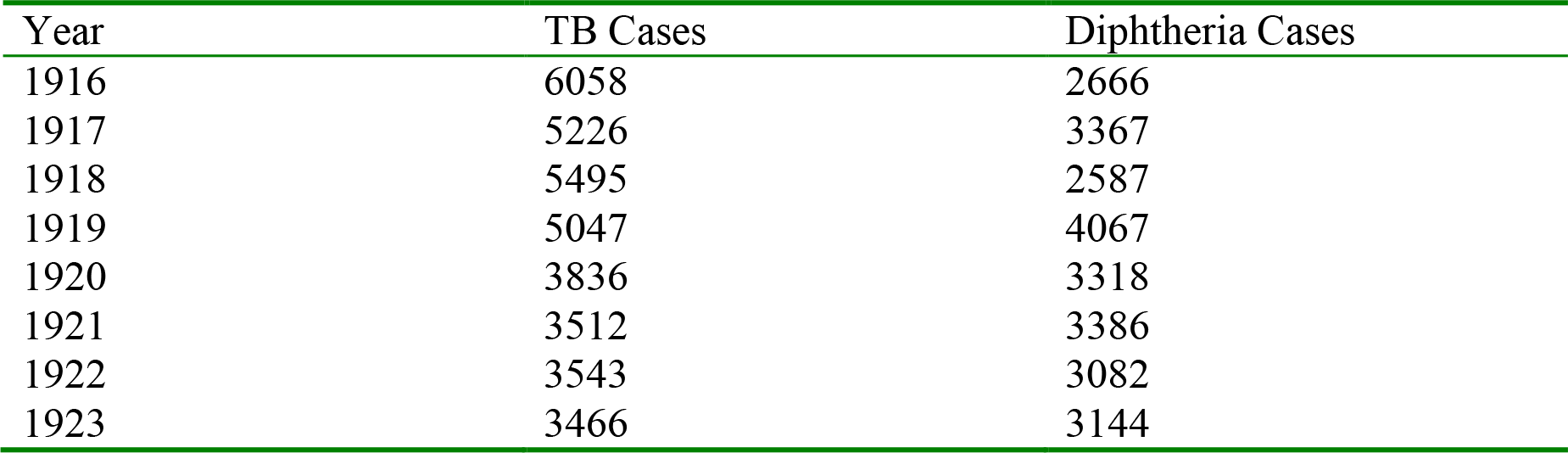
Philadelphia, TB and diphtheria cases, 1916-1923.

Regression models for each city have about the same strength of fit as measured by R^2^ (Table 6), ranging from 0.51 in Boston to 0.68 in Detroit. Figures 3-7 shows the estimated model plotted against the actual weekly diphtheria values for each city from 1916 to 1923 (from 1915 for Detroit). Most of the explained variation is accounted for by the polynomial and dummy terms, which capture the strong annual and weekly changes in diphtheria. Further analysis also showed that there was a large diphtheria epidemic in New York in 1921 continuing into early 1922, which made the model less successful. In fact, 1921 had the highest recorded number of diphtheria cases in American history. So New York was re-estimated without 1921 data; both results are in Table 6. The coefficient for tuberculosis is statistically significant in all the cities, with 1921 excluded in New York. Given the standard errors, estimates of the tuberculosis coefficient are most accurate in Boston, Chicago and Detroit, where it varies from 0.11 to 0.135—a very narrow range; the difference among these values is not statistically significant. The Chicago estimate at 0.11 is likely the most accurate. Further inspection of the graphs shows that the timing of the annual peaks of diphtheria cases in Philadelphia and, especially, in New York City (Fig. 6) are more irregular, less seasonal, than in the other cities. The reason for this is unknown, but it reduces the success of the curve-fitting model and, possibly, makes it harder to detect a consistent relationship between diphtheria and tuberculosis.

**Figure 3.**
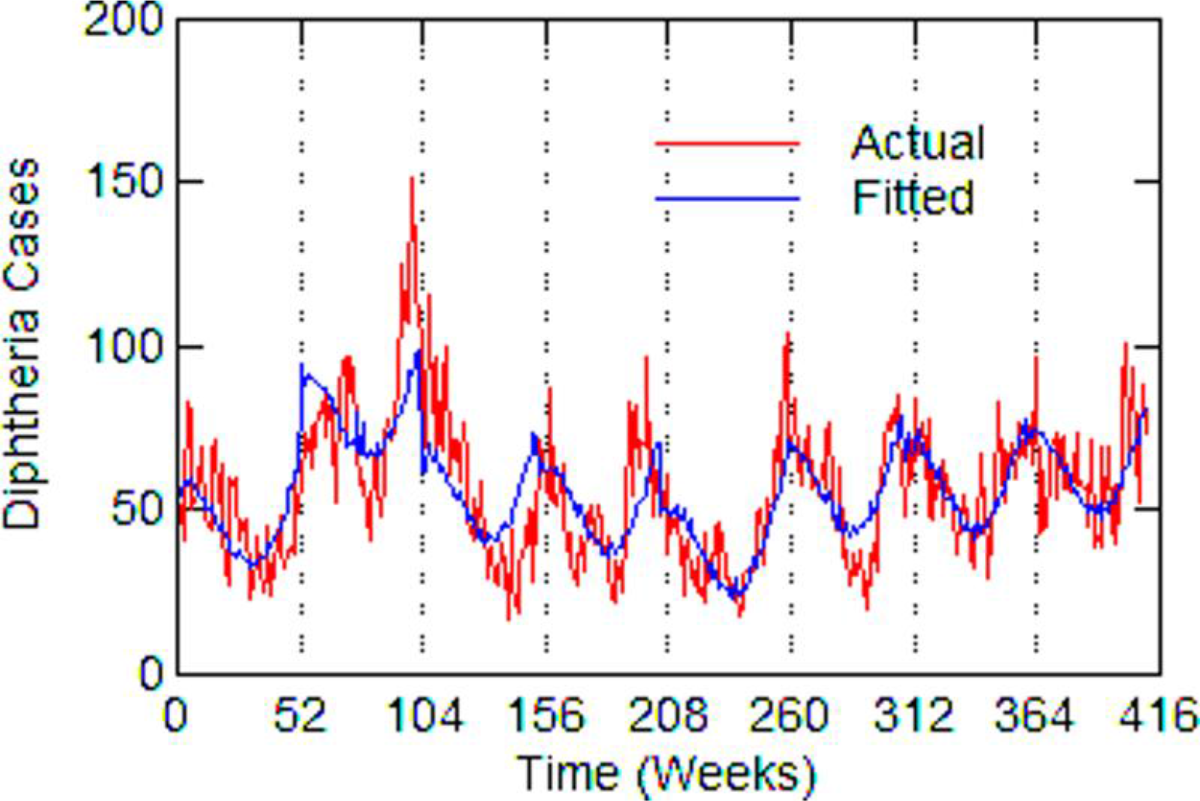
Boston, actual and fitted (estimated) diphtheria cases by week from 1916 to 1923.

**Figure 4.**
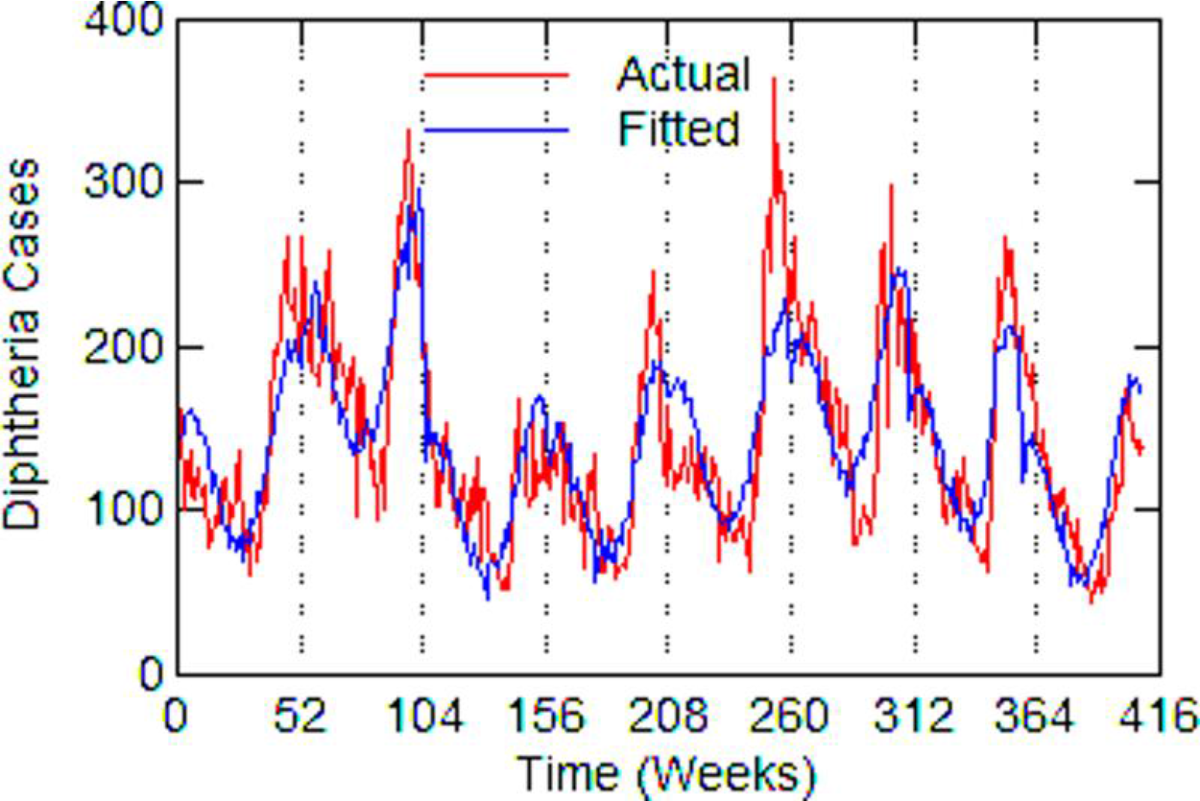
Chicago, actual and fitted (estimated) diphtheria cases by week from 1916 to 1923.

**Figure 5.**
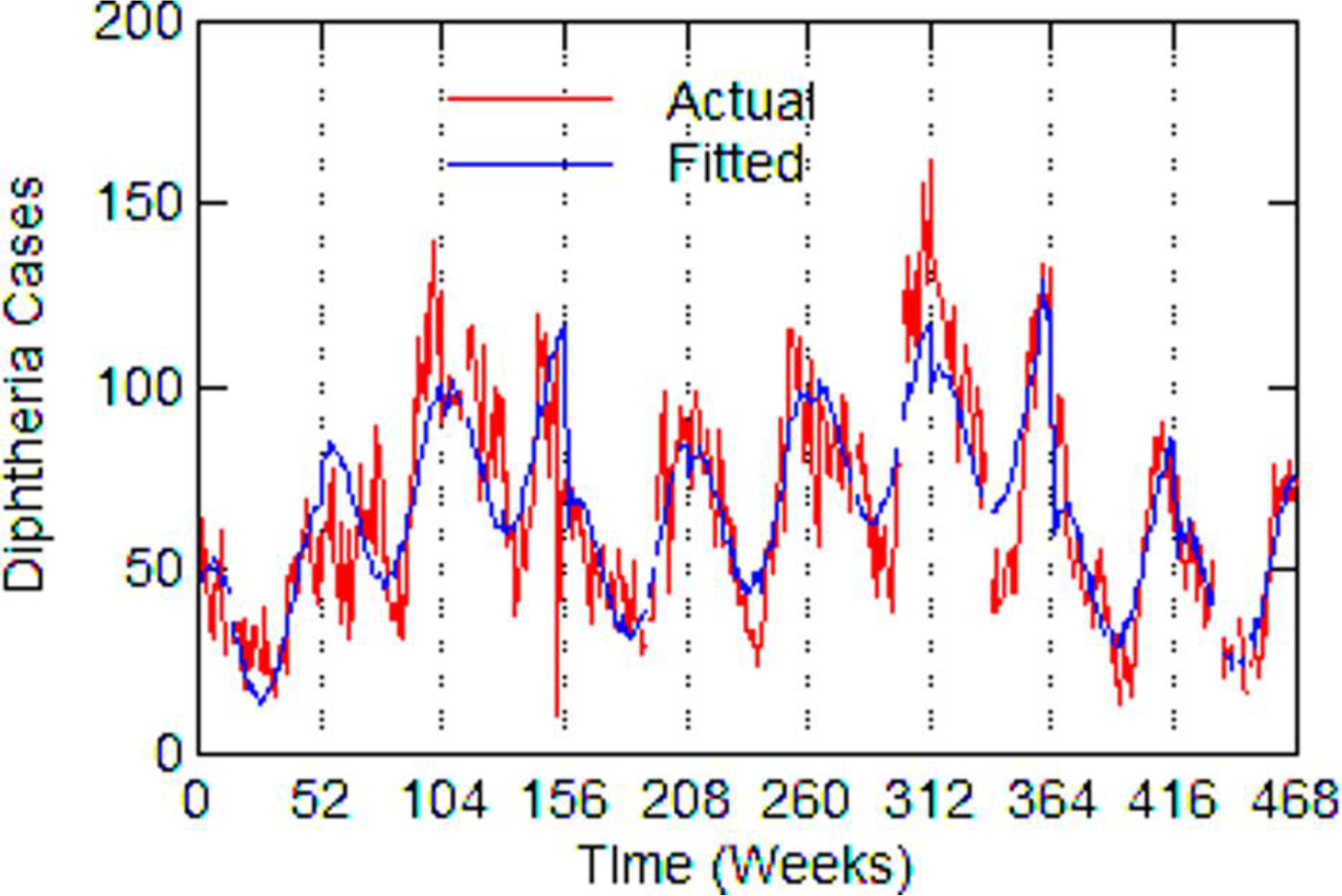
Detroit, actual and fitted (estimated) diphtheria cases by week from 1915 to 1923.

**Figure 6.**
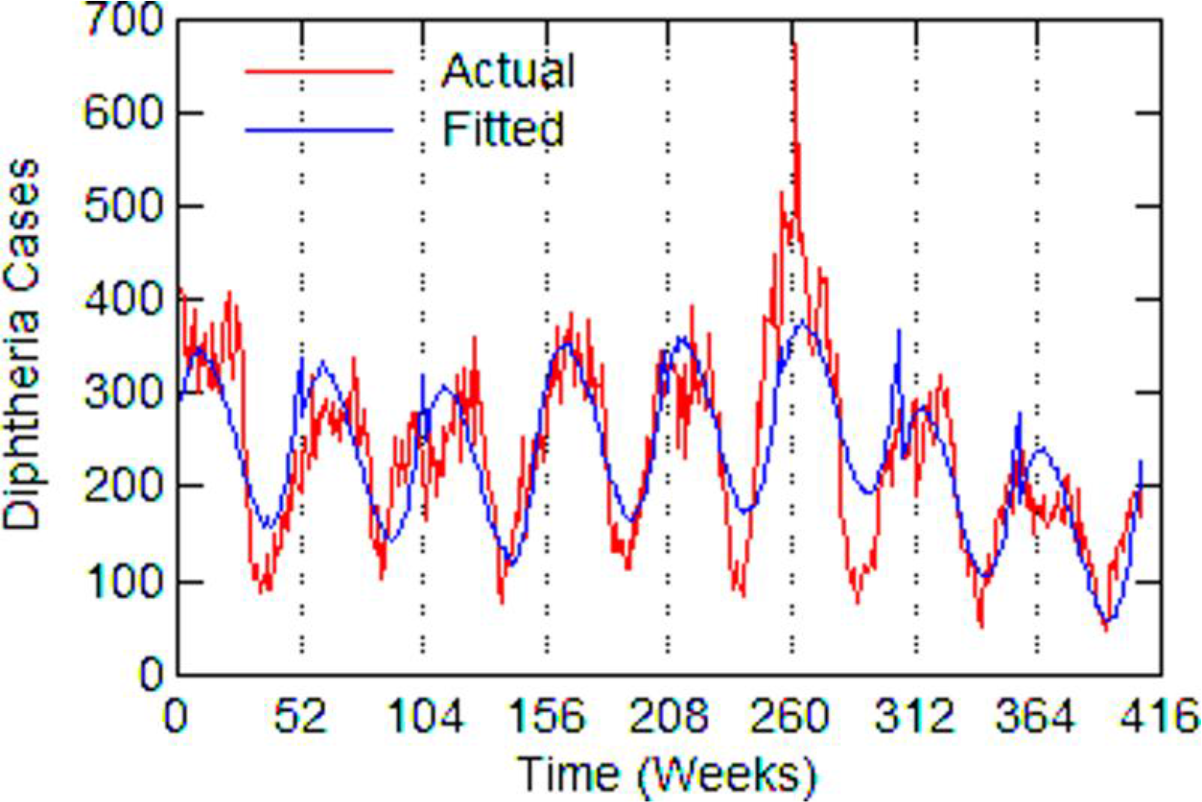
New York City, actual and fitted (estimated) diphtheria cases by week from 1915 to 1923.

**Figure 7.**
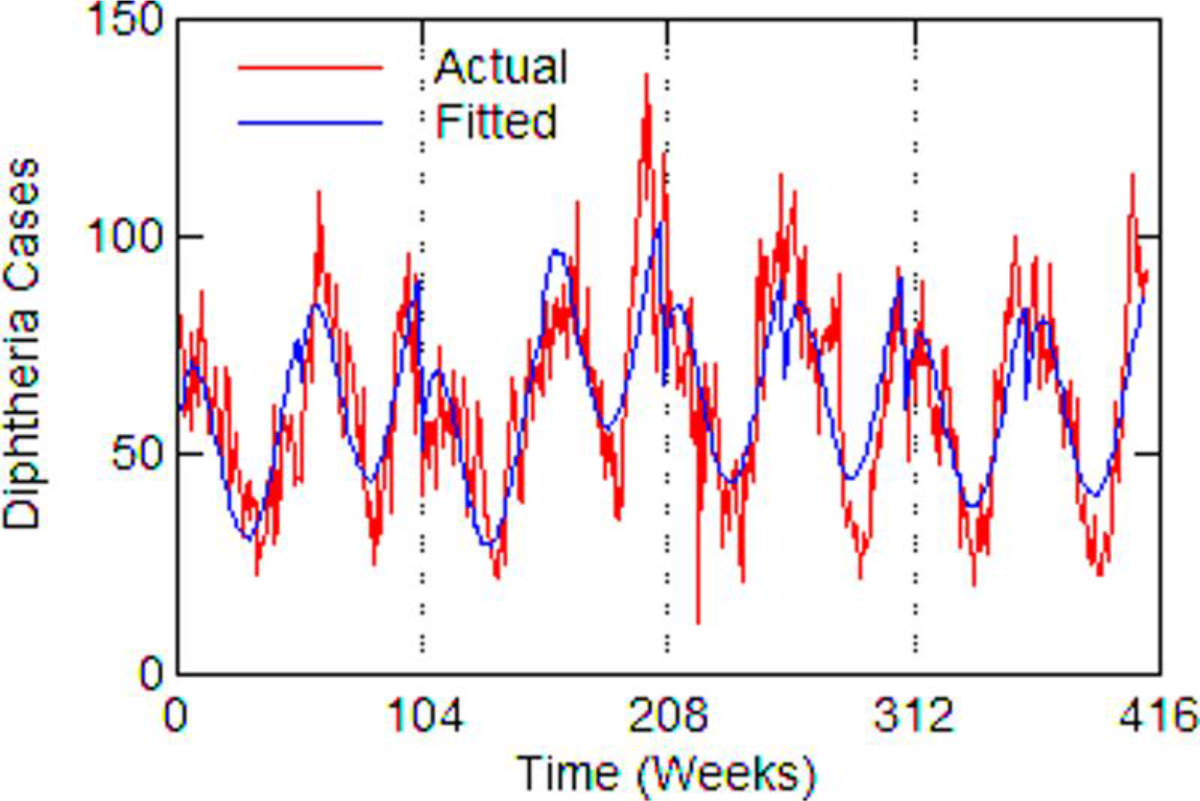
Philadelphia, actual and fitted (estimated) diphtheria cases by week from 1916 to 1923.

**Table 6.**
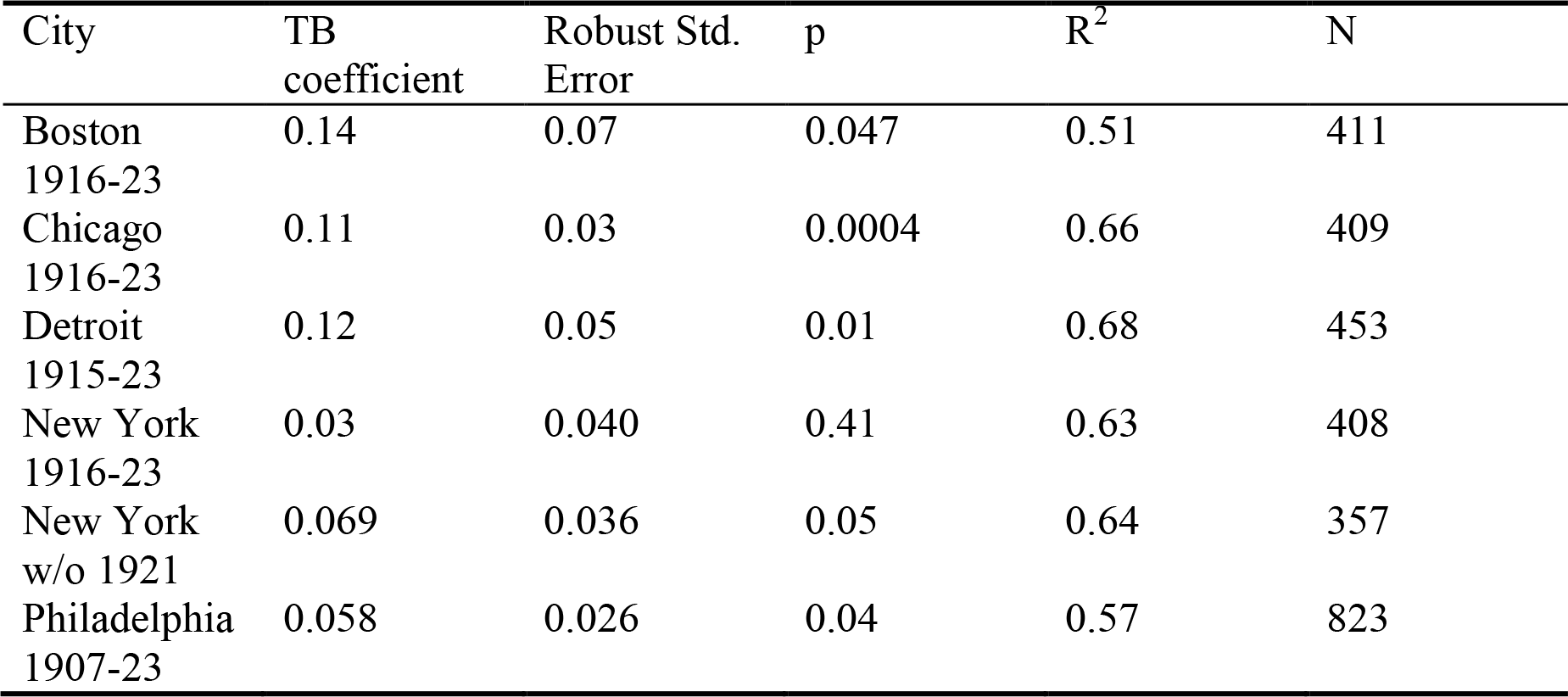
Estimated regression coefficient for tuberculosis in the diphtheria time-series model; polynomial and dummy variable coefficients not reported.

With Chicago as the best case estimate, one can work out roughly how many diphtheria cases might have been associated with tuberculosis. In 1920, in the middle of the time series and after the influenza epidemic of 1918/1919, there were 10,684 reported tuberculosis cases. For a coefficient value of 0.11 with 95% CI [0.05, 0.17], this would mean that about 1,175 [534 – 1,816] diphtheria cases or 15% [7% - 23%] of the total 7,741 diphtheria cases had involved tuberculosis that year. This percentage is similar to rates of association reported in the historical literature.

## Discussion

The statistical analysis supports the historical reports within the inherent limitations of aggregate data analysis, which is that one does not know how strongly the findings apply at the individual level. But it is the best one can do with historical public health data. In any case, the results of both the historical review and statistical analysis call on us to consider possible explanations for a connection between tuberculosis and diphtheria. The medical literature does not offer an immediate answer for this.

Several alternative explanations come to mind. First is the fact that both diseases are fostered by poverty, malnourishment, and overcrowding. These factors may explain differences between cities or how incidence in a city may change over a long time period, but these factors are unlikely to affect short-term or week-to-week variation in disease incidence. Another possibility is the spread of disease by contaminated milk before pasteurization. Milk-borne diseases included diphtheria, scarlet fever and strep throat, tuberculosis and bovine tuberculosis, and typhoid fever.[11] Analysis of milk-borne diseases in Massachusetts from 1909 to 1913 concluded, however, that transmission of diphtheria through milk was negligible.

One must look more closely at the cell biology of tuberculosis and diphtheria for clues to their association. As different as the two diseases are, both bacteria have the same type of protective cell wall—an extra layer of fatty cells--which heightens pathogenicity.[12,13] This may give a synergistic benefit to both diseases in a co-infection, and it is also a potential target for drug development against both.[14]

A better clue to the association of tuberculosis and diphtheria may be how the bacteria obtain and use iron. Bacteria need iron to grow, although the amount of iron must be carefully controlled for survival. The body defends itself against disease by strategies to withhold iron.[15] Tuberculosis is highly dependent on the availability of iron in the body and is so effective at obtaining iron that it can cause anemia; but a person with iron-deprivation anemia has greater resistance to tuberculosis.[16,17]

Diphtheria has a more complex response to iron. Iron activates a gene that represses the production of diphtheria toxin and other components of its iron acquisition system making the disease less virulent, whereas a low level of extracellular iron causes diphtheria to release its dangerous toxin.[18,19] Because of the effect of low iron levels on the virulence of diphtheria, one might suspect that diphtheria could be treated with an iron-based therapy. And, in fact, in the late 1800s compounds of iron, such as ferric subsulphate (Monsel’s styptic), were used as a topical treatment for diphtheria.[20] It is now known that tuberculosis and diphtheria have the same, unique type of iron-dependent repressor genes, which control the assimilation of iron.[21] With this information, one can consider the possible result of a co-infection. Tuberculosis, by extracting available iron in the body, would enhance the production of diphtheria toxin, substantially worsening the outcome of that disease. Debilitation caused by a severe diphtheria infection might then exacerbate or reactivate a preexisting tuberculosis infection. Whether this actually happens, however, is matter for future research.

## Conclusion

This research revives historical medical information about the relationship between diphtheria and tuberculosis, while adding a statistical analysis that would not have been possible in an earlier era. Past medical research has not given a specific cause for the relationship between diphtheria and tuberculosis. However, contemporary research on their bacterial cell walls, iron assimilation, and iron-dependent repressor genes point toward possible bases for their association.

Although one might think of diphtheria epidemics as long-ago events, this is not necessarily true. Tuberculosis is still widespread in many parts of the world, and diphtheria remains a threat. After the collapse of the Soviet Union in the 1990s, there was a resurgence of several epidemic diseases including the first large-scale diphtheria epidemic in over three decades with over 140,000 cases.[22] The combination of a failure to vaccinate children and susceptible adults quickly brought diphtheria back. But, as in the historical period, the resurgence of diphtheria may have been accelerated because tuberculosis is widespread in Russia and often drug resistant. By the end of the 1990s there were over 300,000 new tuberculosis cases, a million or more recovered patients, and as many again with positive tuberculin tests.[23] Further research on the possible relationship between diphtheria and tuberculosis in Russia is warranted.

## Competing or Conflicting Interests

The author has no competing interests.

## Ethical Statement

This research used only aggregate, historical, statistical public health data that is in the public domain. No human subjects were involved.

## Data Accessibility

This data is available to the public at no cost from Project Tycho at the University of Pittsburgh (https://www.tycho.pitt.edu/). The city-level data used here is “Level 2” data available at https://www.tycho.pitt.edu/data/level2.php in downloadable Excel files.

